# Salience Network Segregation and Symptom Profiles in Psychosis Prodrome Subgroups

**DOI:** 10.1101/2024.12.06.627190

**Authors:** Aditya Iyer, William Stanford, Eran Dayan, Rose Mary Xavier

## Abstract

**Background:** Understanding neurobiological similarities between individuals with prodromal psychosis symptoms can improve early identification and intervention strategies. Here, we aimed to (i) identify neurobiologically similar groups by integrating resting-state functional connectivity and prodromal symptom data and (ii) discern discriminating symptom profiles and brain connectivity patterns in the identified sub-groups.

**Methods:** Our sample (*N*=922) was extracted from the Philadelphia Neurodevelopmental Cohort and included individuals ages 12-21 years with fMRI and self-reported psychopathology data. Analyses were conducted separately for youth and early adults. We first constructed a two-layer network using pair-wise similarity distances between participants based on resting-state functional connectivity and prodromal positive psychosis symptoms. We then performed community detection via a multiplex stochastic block model to identify subject clusters.

**Results:** We identified 2 blocks or communities for both the youth (*n*=458 and 179) and early adult (*n*=173 and 112) groups. Connection parameter estimates of the neuroimaging layer were nearly identical between blocks for both age groups whereas there was significant variation for the symptom layer. Psychopathology symptom and brain system segregation profiles were consistent across age groups. The youth block (*n*=458) with higher salience network segregation values had higher mean scores for prodromal symptoms. However, the early adult block (*n*=173) with lower salience network segregation had higher mean prodromal scores.

**Conclusions:** By integrating global similarities in brain connectivity and prodromal symptoms, we identified distinct subgroups. These groups show differences in symptom profiles and network segregation in youth and early adults, indicating significant variations in developmental paths for psychosis spectrum.

## Introduction

The psychosis prodrome, a period that can range from a few weeks to several years, precedes the full onset of schizophrenia (1). Several attempts have been made to predict the prodromal period and the prognosis of schizophrenia in high-risk individuals leveraging models incorporating clinical, environmental, or neurocognitive factors (2–4). In studies that examine the neurobiological underpinnings of schizophrenia, patients with schizophrenia exhibit structural deficits in gray matter regions, with cortical thinning in the frontal, temporal, anterior cingulate, and insular cortices (4–6). These regions either comprise the salience network, which is involved in the selection of relevant stimuli (7), or they play a significant role in facilitating and modulating the network’s activation and connectivity. Structural deficits in these regions have also been observed in individuals with high risk for developing psychosis (5,6). However, such deficits are not specific to schizophrenia and can be indicative of other neuropsychiatric disorders (5).

Functional neuroimaging studies of schizophrenia have revealed abnormal connectivity patterns within the salience network and its closely associated regions, including those that comprise the default mode network (DMN), frontoparietal network (FPN), and dorsal attention network (DAN; 5,6,8–13). In particular, anti-correlation between the DMN and the salience network was found to be reduced (10,13), which can be indicative of the confusion of internally and externally focused states and the disruption of cognition—a hallmark of psychotic disorders (13).

Studies have employed network analyses to investigate psychopathology symptoms (14–17). Graph-based models from such studies consist of nodes that represent psychopathological variables and edges that represent the conditional dependencies between two given variables.

These models are effective at specifying symptoms that convey the highest level of clinical information, and they can explain phenomena such as psychiatric comorbidities through the topology of their networks (18). However, they do not identify psychiatric subtypes or participant groups that are most at risk of psychopathology.

Community detection is a possible approach for discovering psychiatric subtypes within larger participant groups. In this approach, clustering is performed on network (or graph) structures with nodes that represent individual participants and edges that represent a measure of similarity between a given subject pair. Compared to traditional clustering methods, community detection has not received as much attention in psychiatric literature (19). Existing work using community detection has observed that early onset schizophrenia patients lack a default mode intrinsic connectivity network present in age matched controls (20), but the small sample size of the study (n=26) limited the conclusions that could be drawn from their results (20,21).

The goal of our study was to identify neurobiologically similar participant groups by integrating data spanning resting state functional connectivity and prodromal psychosis symptoms. We further aimed to discern the discriminating symptom profiles and resting state brain connectivity patterns in the communities we identified. To mitigate the sample size constraints encountered in several neuroimaging studies (21), we leveraged the Philadelphia Neurodevelopmental Cohort (PNC) as our subject pool; the PNC dataset contains functional magnetic resonance imaging (fMRI) and psychopathology history data from over 1,000 youth participants (22).

## Methods and Materials

### Participants

The PNC consists of adolescents and early adults (23). Since late adolescence is a critical period in brain development, it is particularly vulnerable for the onset of psychosis and other psychopathology symptoms (24,25). To better encapsulate this crucial phase in our subject network, we conducted our analyses separately for youth (ages 12–17) and early adults (ages 18– 21). We obtained the PNC data from the database of Genotypes and Phenotypes (dbGaP) after required data access and IRB approvals. This study was exempt approved by UNC IRB #19-1935. The data obtained included information from 9,498 participants ages 8–21 who underwent a detailed cognitive and psychiatric assessment (23). At enrollment, 1,445 of these participants also underwent multimodal neuroimaging (22). Our sample (*N*=1158) consists of participants ages 12–21, for whom subject informant data was acquired. Participants ages 8–11 were excluded from our sample as only collateral informant data was acquired (23). We partitioned our data into two subsets: a youth sample consisting of 833 participants ages 12–17 and an early adult sample consisting of the remaining 325 participants ages 18–21 years. For our study, we integrated resting-state functional magnetic resonance imaging (fMRI; neuroimaging layer) and prodromal positive psychosis symptoms (symptom layer) into a multi-layer network to identify communities that were similar.

Demographic and psychopathology symptoms in the PNC were assessed using GOASSESS, a structured computerized instrument developed from a modified version of the Kiddie-Schedule for Affective Disorders and Schizophrenia (23). Psychosis prodromal symptoms were measured using the ordinally structured, 12 item revised PRIME screen which measures positive sub-psychosis symptoms on a 7-point scale ranging from “0” (definitely disagree) to “6” (definitely agree; 27). Psychopathology symptom data consists of 115 individual item-level responses for 16 major psychopathology domains; response options were binary: either positive (the participant answered “yes”) or negative (the participant answered “no”; 28).

We excluded participants with missing data for the prodromal items, contributing to a final sample size of 922 (637 youth subjects and 285 early adult subjects). This is approximately a 20.4% reduction from our initial sample size. Table 1 presents demographic summary statistics for the final sample, including a breakdown of the youth and early adult subsets. There were 87 participants with psychopathology symptom responses that were reported as unknown in our final sample. All unknown responses belonged to one of the 15 domains with binary response values, with none belonging to the psychosis prodromal symptom items. We assumed each unknown response to be negative; this was shown to have minimal effect on validity (28).

**Table 1.**
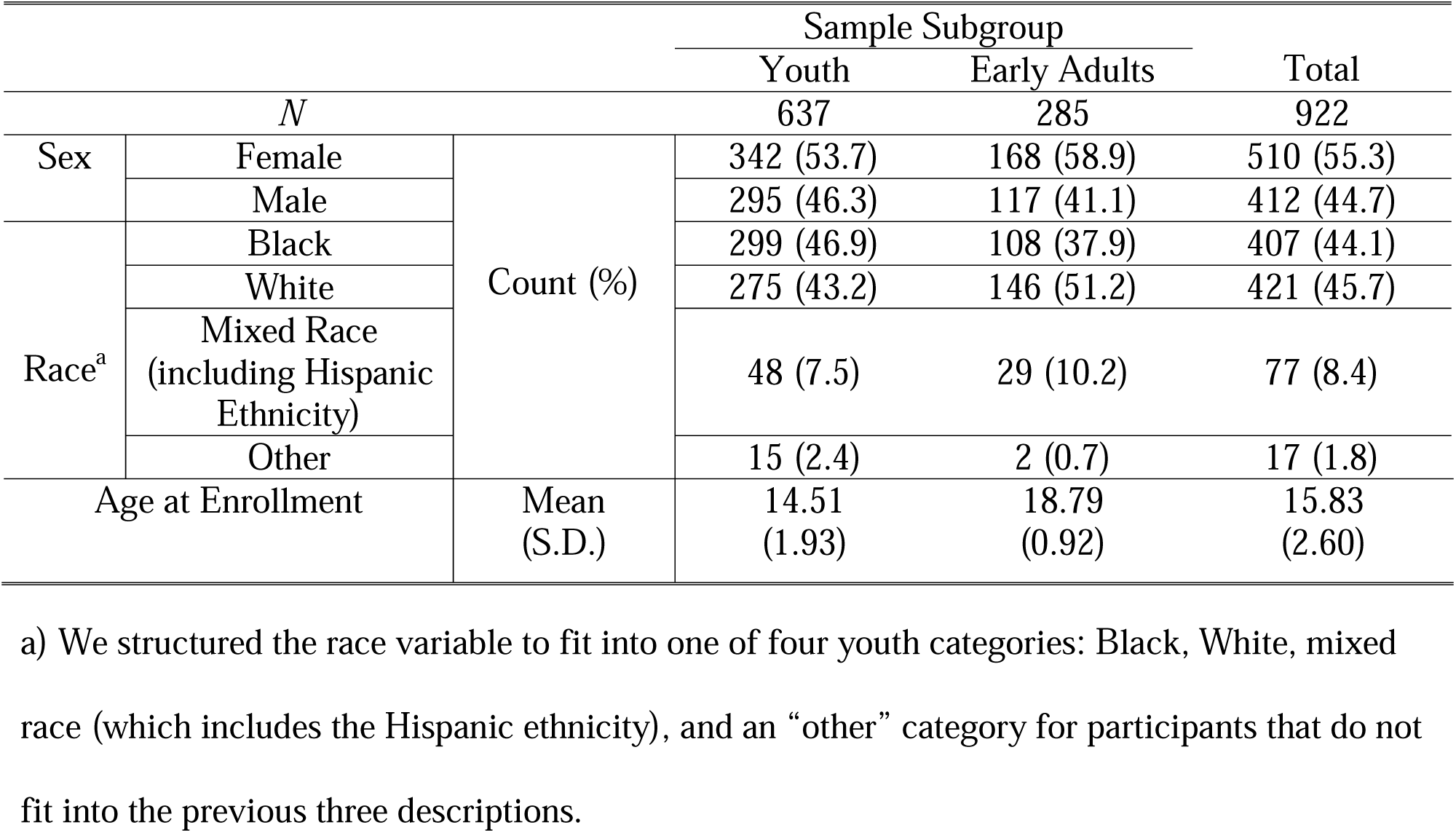
Demographic Summary Statistics of Final Sample (*N*=922)

### Image Acquisition and Preprocessing

Imaging data for PNC subjects were acquired on a 3 Tesla Siemens TIM Trio whole-body scanner. The image acquisition parameters are described elsewhere (22). We preprocessed the neuroimaging data obtained from dbGAP using the CONN toolbox (CONN 21a) running on MATLAB version R2018a (29). Structural images underwent segmentation into gray matter, white matter, and cerebrospinal fluid. Preprocessing of the functional images included realignment and unwarping, slice-timing correction, co-registration to structural images, spatial normalization, and motion outlier identification. White matter, cerebrospinal fluid, ART-based scrubbing, six realignment parameters, and the experimental conditions were included as confound regressors. A temporal band-pass filter was used to remove BOLD frequencies below 0.01 Hz or above 0.1 Hz. Outlier volumes were defined as having greater movement than 0.9 mm or a global signal z-score greater than 3.0. We excluded subjects with less than 80% of volumes removed from subsequent analyses, resulting in a filtered sample of 926 subjects (640 youth and 286 early adults).

### Matrix Construction

Functional time series were obtained using the Gordon parcellation. The cerebral cortex was divided into 333 functionally defined regions of interest (ROIs), which were then partitioned into multiple resting-state networks (30). Weighted FC matrices were constructed for each subject, with edges representing the Fisher *z*-transformed correlations between the functional time series for each ROI.

### Multiplex Network Construction

We constructed a two-layer multiplex network based on pair-wise similarity distances between subjects. The two possible links (connections) between subjects are their similarities in resting-state FC (neuroimaging layer) and responses to psychosis prodromal symptom assessment (symptom layer); altered coordination between the salience network and other brain regions may be associated with prodromal symptoms (10). The construction of each network, including descriptions of the distance measures used, is detailed in the Supplementary Methods. Separate networks were constructed for the youth and early adult subgroups.

### Community Detection

We fit a multiplex stochastic block model (SBM) to each multi-layer network. The SBM is a generative model used to describe the structure of random graphs, finding practical use in community detection (31). It assumes that nodes in a network can be partitioned into multiple blocks (communities). Unlike other commonly used community detection techniques, such as hierarchical clustering and modularity optimization methods, the SBM provides probability distributions to parameterize the connections (edge weights) between and within each community. A formulation of the multiplex SBM and its estimation process are described in the Supplementary Methods.

### Statistical Analyses

We used the *estimateMultiplexSBM* function from the *sbm* R package to fit our models (32), and we subsequently analyzed the participant groups obtained from this procedure. All analyses were performed using Python (v3.9.7) and Jupyter Notebooks (v6.4.5) on the Longleaf computer cluster at the University of North Carolina at Chapel Hill. The NumPy (v1.22.4), pandas (v1.3.4), SciPy (v1.7.1), and scikit-learn (v0.24.2) libraries were used for data preprocessing and statistical computation. The Matplotlib (v3.8.1) and seaborn (v0.11.2) libraries were used for data visualization.

### Brain system segregation

A measure of system segregation, a promising biomarker for psychopathology, was computed to examine the extent to which different cognitive processes are localized to distinct brain networks (33–35). In line with previous implementations (36,37), we measured segregation as the difference between mean within-system (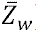) and mean between-system (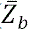) connectivity divided by mean within-system connectivity:

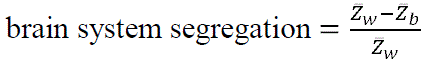

As such, a high system segregation value indicates that the brain networks each tend to partake in unique and specialized functions (35). Conversely, a low system segregation value suggests that the networks are functionally integrated, often partaking in similar tasks or tasks that are interdependent upon another (35). We calculated 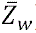 and 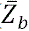 using correlations from participants’ FC matrices. We excluded negative functional connectivity values by setting them to zero, as this has been shown to improve the reliability of graph measures (38). Segregation was evaluated for all participants in both age groups. We computed pairwise measures between the salience network, DMN, FPN, and DAN. These networks are closely linked, and abnormal segregation between the salience network and the other three networks is associated with psychosis symptoms (10,39). We also computed segregation values for each of the four networks individually such that 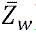 represents the mean of all pairwise edges (correlations) between nodes (ROIs) of the same network and 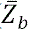 represents the mean of all edges between nodes of the respective network and all other nodes.

### Psychopathology Symptom Analyses

Responses to prodromal symptoms were averaged for each subject. Cross validation of the clustering was conducted using psychopathology symptom responses. Since psychiatric comorbidities are common in patients with psychosis/schizophrenia, successful clustering of the participants should yield significant differences in (non-prodromal) psychopathology symptoms between blocks (40). Supplementary Table S1 presents mean psychopathology symptom scores in the study sample. We examined block-wise means to identify any differences in symptom scores using appropriate statistical tests (Tables S2 and S3).

## Results

### Youth Sample

#### Block Demographics

Table 2 presents demographic summary statistics broken down by block. Distributions of race and gender across age groups and blocks are presented in Figure S1. Youth block 1 contains considerably more Black participants than White participants, but this is reversed in block 2. We found a significant association between gender, age group (youth and early adults), and community (*X*^2^ =15.96, *p*=1.15×10^−3^; Figure S1). Among youth, associations between gender and community (*X*^2^=3.38, *p*=6.61×10^−2^; Figure S1) and race and community were not significant (*X*^2^=3.26, *p*=3.53×10^−1^).

**Table 2.** Demographic Summary Statistics for Each Block (*N*=922)

|  |  |  | Sample Subgroup |  |  |  |
| --- | --- | --- | --- | --- | --- | --- |
|  |  |  | Youth |  | Early Adults |  |
| Block |  |  | 1 | 2 | 1 | 2 |
| *n* |  |  | 458 | 179 | 173 | 112 |
| Sex | Female | Count (%) | 235 (51.3) | 107 (59.8) | 98 (56.6) | 70 (62.5) |
|  | Male |  | 223 (48.7) | 72 (40.2) | 75 (43.4) | 42 (37.5) |
| Race <sup>a</sup> | Black |  | 225 (49.1) | 74 (41.3) | 74 (42.8) | 34 (30.4) |
|  | White |  | 189 (41.3) | 86 (48.0) | 83 (48.0) | 63 (56.3) |
|  | Mixed Race<br>(including<br>Hispanic<br>Ethnicity) |  | 34 (7.4) | 14 (7.8) | 15 (8.7) | 14 (12.5) |
|  | Other |  | 10 (2.2) | 5 (2.8) | 1 (0.6) | 1 (0.9) |
| Age at Enrollment |  | Mean<br>(S.D.) | 14.45<br>(1.95) | 14.64<br>(1.87) | 18.76<br>(0.94) | 18.84<br>(0.90) |

a) We structured the race variable to fit into one of four youth categories: Black, White, mixed race (which includes the Hispanic ethnicity), and an “other” category for participants that do not fit into the previous three descriptions.

#### Parameter Estimates

As presented in Figure 1C, the within-block parameter estimates (means and variances) for the neuroimaging layer are nearly identical for the two youth blocks (*µ* ≈ 0.30 and *σ*^2^ ≈ 0.04). These values are also close to the estimate for the connection parameter between blocks 1 and 2 (*µ* ≈ 0.29 and *σ*^2^ ≈ 0.04). In contrast, there is considerably more variation in the symptom layer, suggesting that the clustering by the variational EM algorithm was influenced more by the symptom layer than the neuroimaging layer.

**Figure 1.**
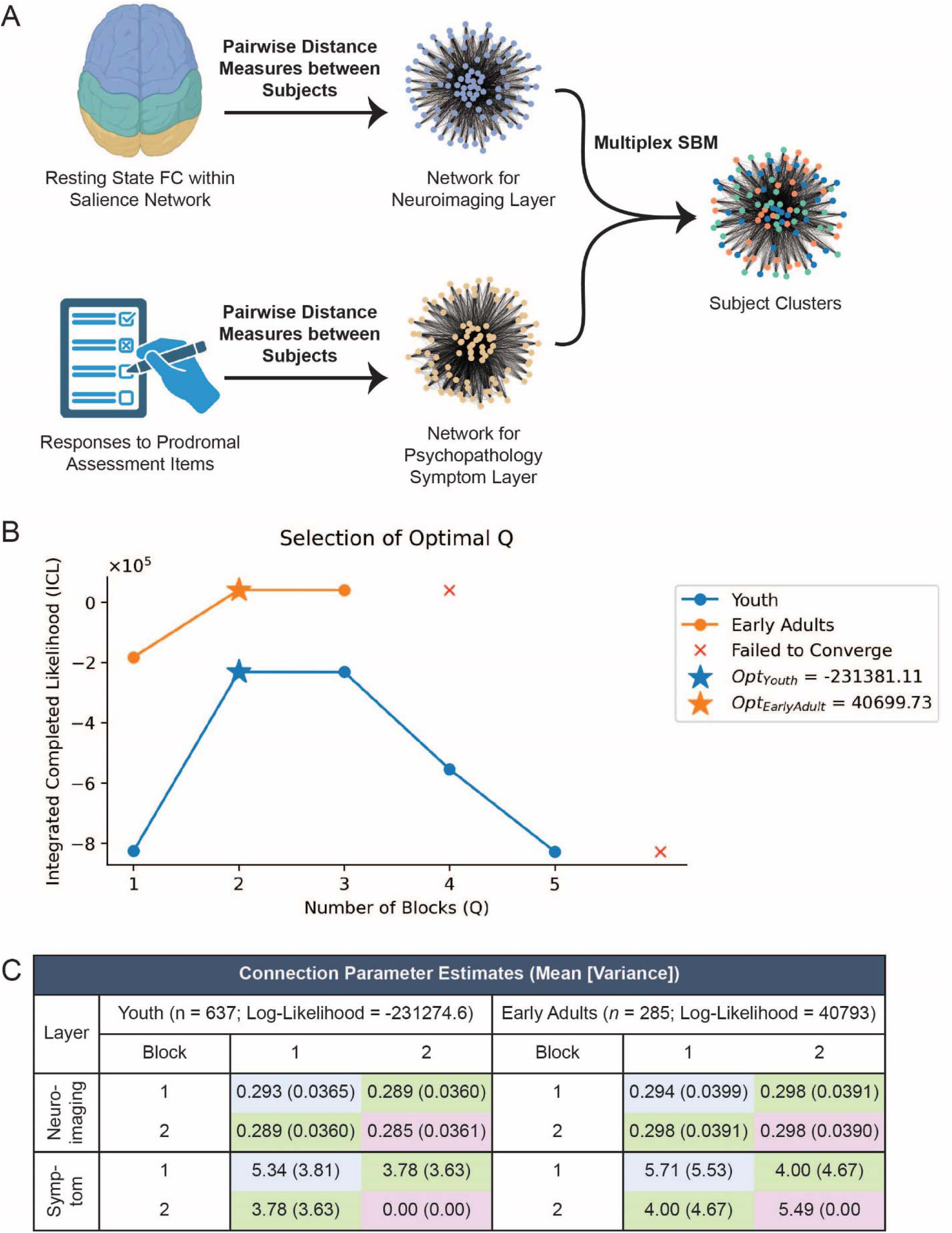
Training of the multiplex stochastic block model (SBM). A diagram of the data pipeline is shown in (**A**). We integrated participant data spanning (1) resting state functional connectivity (FC) within the salience brain network and (2) responses to the prodromal symptom assessment items. Pair-wise similarity distance measures were computed between participants to produce a weighted graph (network) with a neuroimaging layer and a (prodromal) symptom layer. A multiplex SBM was fit to the multi-layer network via a variational expectation-maximization (EM) algorithm. This process was conducted twice, once for the youth and again for the early adults. The selection process for the optimal number of blocks (*Q*) is shown in (**B**). We selected the optimal *Q* for each subject group (youth and early adults) using the integrated completed likelihood (ICL) criterion. We evaluated values of *Q* ranging from one to seven and selected the models that converged with the highest ICL. Convergence was not reached when *Q* was greater than six for the youth and four for the early adults. We found *Q*: = 2 to be optimal for both the youth and early adults, producing ICLs of –231,381.11 and 40,699.73, respectively. A connection parameter estimates table for the multiplex stochastic block models is shown in (**C**). We report the mean and variance of the within– and between-block estimates for each layer in each age group’s model. We also report the log-likelihoods of the models.

#### Brain System Segregation

We evaluated brain system segregation for four different pairings of functional networks: (1) the salience network and all other regions of interest, (2) the DMN and all other regions of interest, (3) the FPN and all other regions of interest, (4) the DAN and all other regions of interest. All mean segregation values were positive (BH-adj *p* < 0.05; Figure 2), indicating that within-network connectivity tended to be stronger than between-network connectivity for the evaluated functional network pairings. Group comparisons of the four measures were significant in both communities (block 1: *F*=31.88, *p*=4.39×10^−20^); block 2: *F*=8.32, *p*=1.90×10^−5^; Figure 2). Segregation for the salience network was the highest (Figure 2). All pairwise post-hoc comparisons (two-tailed *t*-tests) involving the salience network were significant (Benjamini-Hochberg-adjusted [BH-adj] *p* < 0.05) for block 1 but not for block 2 (Figure 2C).

**Figure 2.**
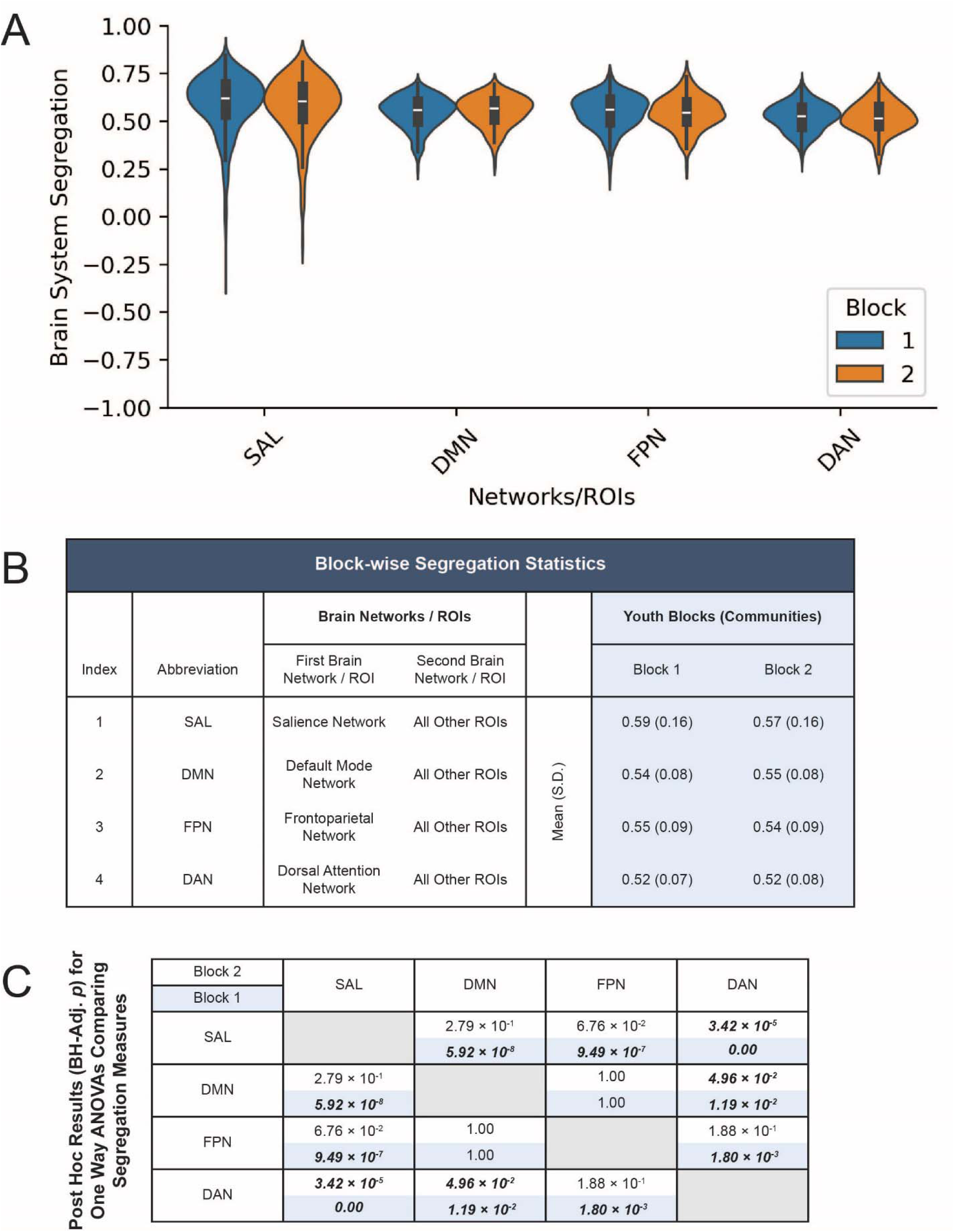
Brain system segregation values for youth participants (*n*=637) broken down by block (Block 1: *n*=458; Block 2: *n*=179). A visualization of the block-wise distributions of segregation values is shown in (**A**). A table with block-wise segregation statistics (means and standard deviations) is shown in (**B**). A symmetric matrix with post-hoc comparison (two-tailed *t*-test) results for one-way ANOVAs that assess differences between the four segregation measures is shown in (**C**). An ANOVA was conducted separately for Block 1 (*F*=31.88, *p*=4.39×10^−20^) and Block 2 (*F*=8.32, *p*=1.90×10^−5^) of the youth. Significant pair-wise *t*-test results (BH-adj. *p*) are bolded and italicized. Significance was determined based on a threshold of 0.05. The Benjamini-Hochberg method was used to adjust for multiple comparisons.

#### Psychopathology Symptoms

For the two youth communities, psychopathology symptoms scores were higher for the attention deficit hyperactive domain (ADD), depression (DEP), the generalized anxiety domain (GAD), mania (MAN), the oppositional defiant domain (ODD), specific phobias (PHB), and the social anxiety domain (SOC; Figure 3A). Of these seven domains, four (DEP, GAD, MAN, and SOC) are known to be closely linked with psychosis (40). Block 1 had higher values for all symptom measures assessed. The two blocks exhibited the strongest differences in ADD (BH-adj. *p*=5.47×10^−13^), MAN (*p*=6.54×10^−11^), and the psychosis domain (PSY; BH-adj. *p*=4.65×10^−11^), but 14 out of the 15 psychopathology domains assessed showed significant differences (BH-adj. *p* < 0.05), with the exception of agoraphobia (Supplementary Table S2).

**Figure 3.**
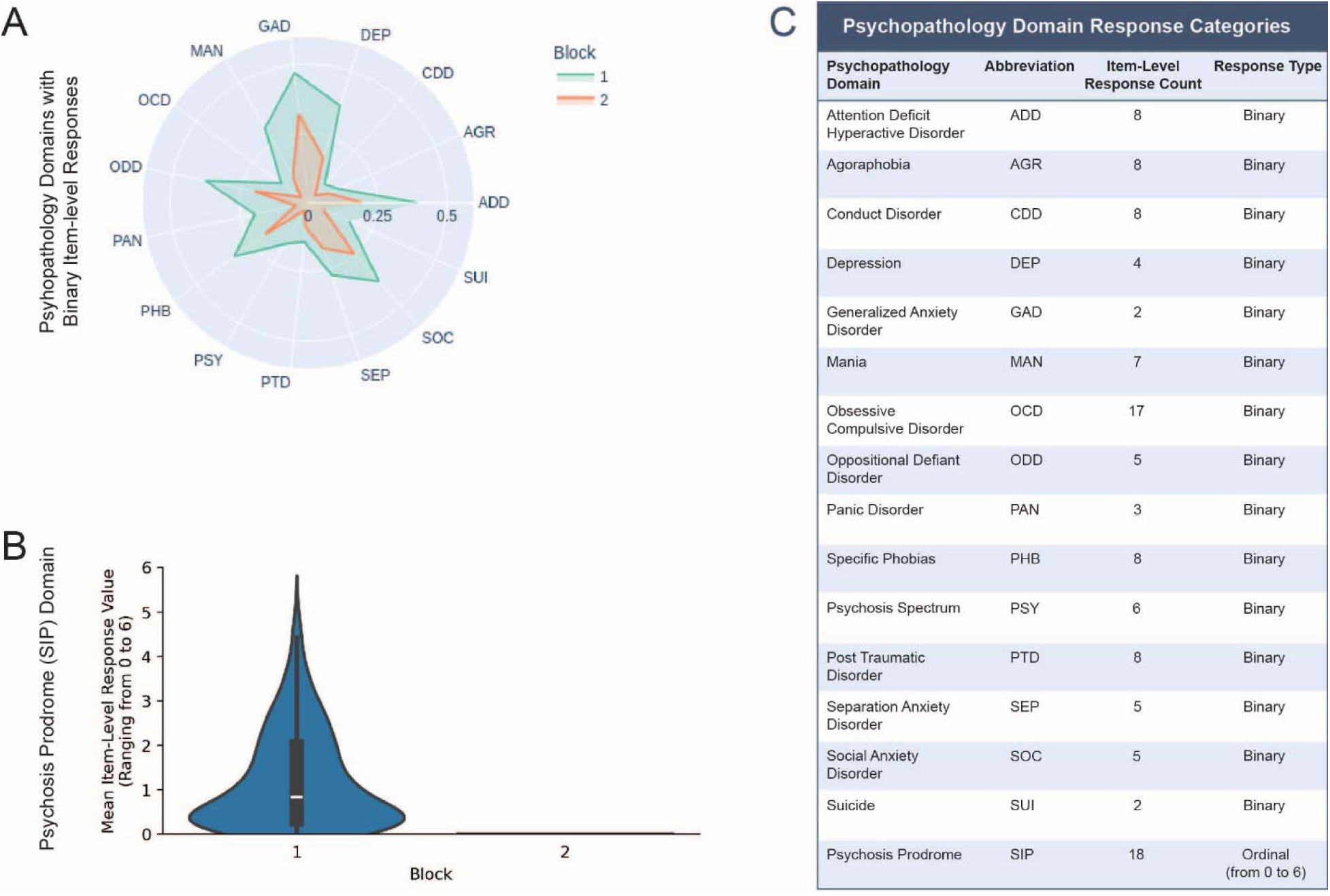
Psychopathology symptom scores for youth participants (*n*=637) broken down by block (Block 1: *n*=458; Block 2: *n*=179). The mean positive (“yes”) response count per subject for each of the 15 non-prodromal psychopathology domains is shown in (**A**). The means are scaled to range from 0 to 1 to better visualize the differences between blocks. The distributions of the mean response values for the prodromal psychosis items are shown in (**B**). A table displaying the abbreviation, number of item-level responses, and response value type (binary or ordinal) for each psychopathology domain is shown in (**C**).

Youth in block 1 (*n*=458) had a mean value of 1.2 (SD = 1.1) for prodromal symptoms (Figure 5B), whereas for block 2 (*n*=179) the mean was 0.0 (SD = 0). Block 1 also had higher salience network segregation than block 2 (Figure 2) and significant post-hoc segregation comparisons involving the salience network (Figure 2C) suggesting an association between prodromal psychosis symptoms and abnormal salience network segregation among youth. Since our community detection model incorporates both functional connectivity measures and psychosis prodrome scores to cluster participants, we did not conduct between-block significance tests involving the segregation measures or prodrome scores; this circular analysis would produce artificially inflated test statistics.

### Early Adult Sample

#### Block Demographics

There is a higher proportion of African Americans in block 1 than block 2 in the early adult group (Table 2; Supplementary Figure S1). However, the association between race and community is not significant (*X*^2^=4.75, *p*=1.91×10^−1^; Supplementary Figure S1). Consistent with youth, we did not find a significant association between gender and community among early adults (*X*^2^=0.74, *p*=3.91×10^−1^; Supplementary Figure S1).

#### Parameter Estimates

Similar to our findings within the youth subgroup, all connection parameters for the neuroimaging layer are nearly identical (*µ* ≈ 0.30 and *σ*^2^ ≈ 0.04; Figure 1C). The early adult connection parameters are also close to those observed in the youth subgroup, suggesting a lack of change in salience network functional activity between youth and early adulthood. Once more, we found considerably more variation in the mean and variance of the symptom layer edge weights.

#### Brain System Segregation

Group comparisons of the four segregation measures were significant in both early adult communities (block 1: *F*=2.89, *p*=3.49×10^−2^; block 2: *F*=5.65, *p*=8.29×10^−4^; Figure 4). The segregation patterns of the early adult communities closely mirror those of the youth communities: salience network segregation tended to be higher than the segregation values for each of the DMN, FPN, and DAN (Figure 4). However, all pairwise post-hoc comparisons (two-tailed *t*-tests) involving the salience network were not significant (BH-adj *p* < 0.05), apart from the comparison between the salience network and DAN in block 2 (BH-adj *p*=4.96×10^−2^; Figure 4C). In fact, only three of the 12 post-hoc comparisons were significant for the early adult communities, a departure from the trend observed among youth (Figure 4C). Given that the early adult subgroup consists of 285 participants, with each of its communities consisting of over 100 participants, the lack of associations for the segregation measures is likely not an artifact of lower statistical power.

**Figure 4.**
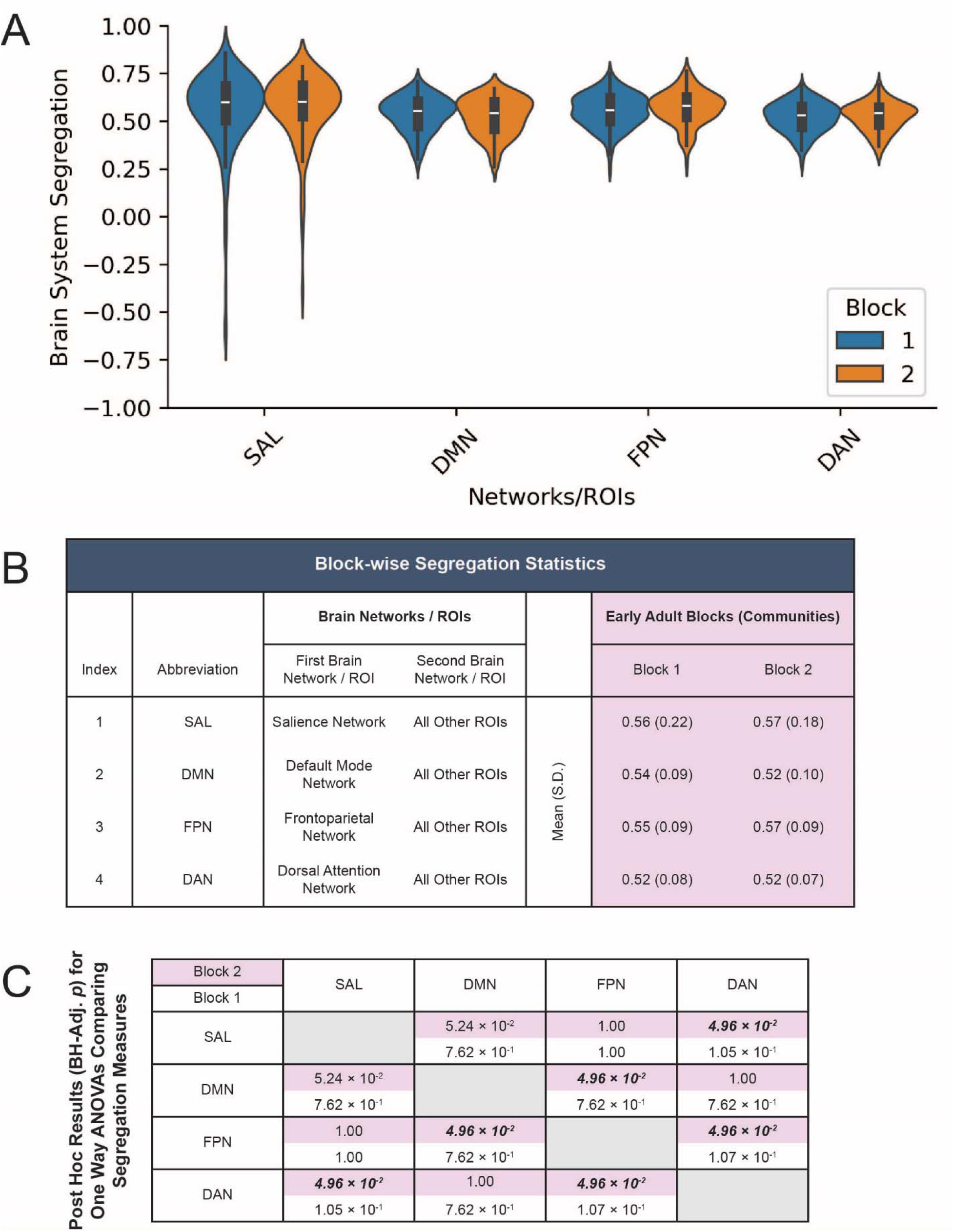
Brain system segregation values for early adult participants (*n*=285) broken down by block (Block 1: *n*=173; Block 2: *n*=112). A visualization of the block-wise distributions of segregation values is shown in (**A**). A table with block-wise segregation statistics (means and standard deviations) is shown in (**B**). A symmetric matrix with post-hoc comparison (two-tailed *t*-test) results for one-way ANOVAs that assess differences between the four segregation measures is shown in (**C**). An ANOVA was conducted separately for Block 1 (*F*=2.89, *p*=3.49×10^−2^) and Block 2 (*F*=5.65, *p*=8.29×10^−4^) of the early adults. Significant pair-wise *t*-test results (BH-adj. *p*) are bolded and italicized. Significance was determined based on a threshold of 0.05. The Benjamini-Hochberg method was used to adjust for multiple comparisons.

Furthermore, consistent with the observation among youth, the four segregation measures have positive means for the early adults (BH-adj *p* < 0.05; Figure 4). From visual examination, the distributions of block-wise segregation measures appear to be similar between youth and early adults. Overall, these similarities indicate a minimal change in salience network functional activity between youth and early adulthood and a lack of alteration in the global functional connectivity signal by community detection for the examined networks.

#### Psychopathology Symptoms

The psychopathology symptom profiles of the early adult communities resemble those of the youth communities, with blocks 1 and 2 showing the strongest differences in ADD (BH-adj. *p*=2.06×10^−5^), MAN (*p*=1.40×10^−5^), and PSY (BH-adj. *p*=2.06×10^−5^). All 15 of the psychopathology domains assessed exhibited significant differences between the two blocks (BH-adj. *p* < 0.05; Supplementary Table S3). As observed among the youth, multiple participants clustered in Block 1 (*n*=173) experienced prodromal symptoms (mean 0.9, SD 0.9; Figure 5B), whereas participants in Block 2 (*n*=112) reported no prodromal symptoms (mean 0, SD 0). However, contrary to the pattern among youth, Block 1 has lower salience network segregation than Block 2 (Figure 4). Additionally, none of the pairwise post-hoc comparisons for block 1 that involve the salience network are significant (Figure 4C). This indicates that, among early adults, the association between prodromal symptoms and salience network segregation disappears.

**Figure 5.**
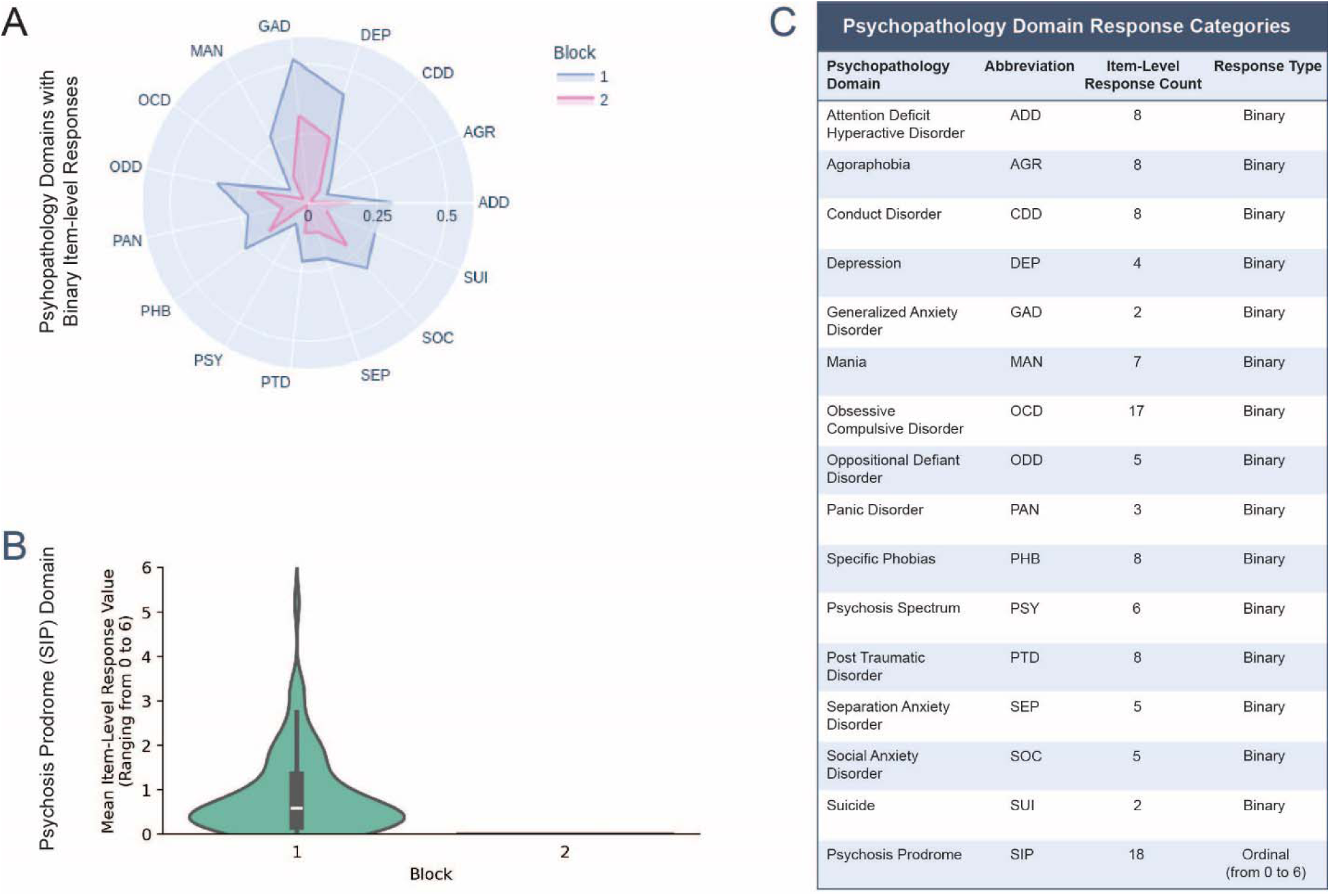
Psychopathology symptom scores for early adult participants (*n*=285) broken down by block (Block 1: *n*=173; Block 2: *n*=112). The mean positive (“yes”) response count per subject for each of the 15 non-prodromal psychopathology domains is shown in (**A**). The means are scaled to range from 0 to 1 to better visualize the differences between blocks. The distributions of the mean response values for the prodromal psychosis items are shown in (**B**). A table displaying the abbreviation, number of item-level responses, and response value type (binary or ordinal) for each psychopathology domain is shown in (**C**).

## Discussion

To identify neurobiologically similar participant sub-groups, we fit multiplex SBMs to two-layer networks constructed using PNC participants’ psychopathology histories and resting state FC within the salience network. Overall, we find that increased salience network segregation is associated with prodromal psychosis symptoms in youth, but not in early adults.

### Psychopathology History is More Influential than Functional Brain Activity for Psychiatric Subtyping

Among both the youth and early adults, community detection was driven mostly by psychopathology history over pair-wise distance measures of resting state functional activity. This imbalance can possibly be attributed to the sources of data used to construct each network layer. fMRI data used to construct the neuroimaging layer is susceptible to several sources of noise; these include subject motion, instrumentation artifacts such as magnetic field fluctuations, and physiological noise such as variations in heart rate and respiration (41). A low SNR can make it challenging to detect true brain activity related changes (41). This presents a challenge for its use in clinical subtyping, as clustering methodologies may fail to identify meaningful changes in functional brain activity between participant groups.

Our model incorporated fMRI data in the form of pairwise distance measures comprising a multigraph layer. These distances were Pearson dissimilarity scores computed using resting state functional connectivity from all ROIs associated with the salience network. Compressing the entirety of a network’s functional activity into a single scalar quantity ranging from zero to one may not adequately encapsulate a present signal. The use of a different distance measure could better embed the fMRI signal. Geodesic distance measures, for example, have found success in improving participant identification from fMRI data (42,43). However, such approaches may impose inaccurate assumptions on underlying structure of the data. It is possible that fMRI may be too global of a biomarker for identifying individual differences relevant to psychopathology.

The stronger influence of psychopathology histories over the clustering also has implications for clinical diagnostics. Though modest effect sizes of neuroimaging markers has been suggested as a limiting factor for reliable classification of individual cases (44), the joint application of psychopathology symptoms and functional imaging can aid in the discovery of prodromal psychosis subtypes as proven in other forms of psychopathology like attention-deficit/hyperactivity disorder (ADHD) and depression (45). Functional imaging in conjunction with psychopathology symptoms can thus serve as a pathognomonic “fingerprint” for clinical diagnosis.

### Consistency of Functional Brain Activity Between Adolescents and Young Adults

Both the youth and early adults have a similar pattern of brain system segregation distributions for the four between-network measures assessed, possibly indicating a lack of change in the specialization of functional brain activity during the transition from adolescence to very early adulthood. It is well documented that brain networks become more segregated during normal adolescent development, more closely resembling the functional activity patterns observed in young adults (46–49). This is particularly the case for networks associated with higher cognitive or emotional functions, such as the salience network, DMN, FPN, and DAN (47–49). Previous studies involving the PNC have detected increases in both brain system segregation and the modularity of structural networks from youth to early adulthood (47,50). However, these studies incorporated participants of all ages (8-22 years) in the analysis without separating the sample into discrete age groups. These studies did not assess differences in sub-populations among participants of a similar age range.

Subgroup identification of the youth and early adult samples underscores differences in functional neurodevelopment between healthy and high risk (HR) subjects. In both age groups, Block 1 contains higher mean scores for prodromal symptoms, indicating a possible HR sub-population. When assessing segregation measures for each of the four major networks, we find that—among youth—the HR block exhibits higher salience network and FPN segregation, lower DMN segregation, and similar DAN segregation compared to the healthy sub-population (Block 2). However, among early adults, the HR block exhibits lower salience network and FPN segregation and higher DMN segregation; the DAN segregation is still similar compared to Block 2. These results suggest different neurodevelopmental trajectories for different subpopulations that are dependent on psychopathology risk. As such, not all networks may become more segregated during maturation from adolescence to adulthood; a given network could become more segregated, or—conversely—more integrated, depending on an individual’s psychiatric subtype. Therefore, positive prodromal symptoms may confound the relationship between brain system segregation and the development from adolescence to early adulthood.

### The Salience Network and Psychosis

Among youth, the block with higher salience network segregation had higher prodromal symptoms. However, among early adults, it was the block with lower salience network segregation that had higher prodromal symptoms. This indicates that the relationship between psychosis risk and salience network segregation varies by age. Aberrant interactions between the salience network and other regions are a hallmark of psychosis (5,6,8–13), and the knowledge of abnormal salience network segregation patterns can guide future efforts at psychiatric subtyping for psychosis prodrome—especially during the early stages of adolescence. Given the wide range of experiences associated with the prodromal stage of psychosis (1,51), there may also be multiple distinct neurobiological patterns with which it may coincide. One such pattern in adolescence may involve heightened salience network segregation, which—coupled with psychopathology symptoms—may serve as a biomarker for a particular prodromal psychosis subtype. The discovery of such a “fingerprint” can help improve the early detection and intervention in schizophrenia spectrum disorders.

Overall, while further investigation may be required to validate the patterns observed in our results, our study adds to the growing body of evidence for the vital role of salience network and the differences in neurodevelopmental trajectories in psychosis. Considering the infrequent use of community detection for clinical subtyping in psychiatric literature (19), this study showcases the potential of community detection for identification of HR sub-populations.

### Limitations and Future Directions

The limitations of our study primarily arise from our use of the PNC as a subject pool. The PNC is a community sample rather than a clinical sample, so it is unclear whether the clustering and subsequently observed block-wise patterns are generalizable to clinical populations. In addition, we were unable to perform analyses on participants ages 8–11 due to the lack of self-reported symptoms. We did not have access to psychiatric diagnoses for any of the participants. Access to such data can help refine community detection by discerning HR sub-populations based on positive clinical diagnoses. Without access to these ground-truth labels, we used responses to non-prodromal assessment items for validation of our community detection.

This is not an ideal approach, as psychiatric comorbidities are not always present in HR individuals, and it is possible that participants may have not disclosed certain symptoms due to stigma. Lastly, our investigation did not evaluate genetics as a risk factor for psychopathology symptoms. Genomic variables of psychosis risk can potentially be used to construct a third layer in our multiplex network to further delineate biological subtypes of prodromal psychosis as well as psychiatric comorbidities, as psychiatric disorders manifest along genetic continua and share common sources of genetic risk (52).

### Conclusions

Our study subtyped PNC participants using an approach used sparsely in psychiatric literature. The results offer insights into the joint use of psychopathology history and functional brain activity in the identification of psychiatrically at-risk youth. Our findings also add nuance to the changes in functional activity from adolescence to early adulthood while further validating the cardinal role of the salience brain network in psychosis.

## Supporting information

Supremental Information

## Acknowledgements

Our appreciation goes to Dr. Jason Xu from the Statistics Department at Duke University for his invaluable consultations on the statistical models used in this research.

This project was sponsored by The Rockefeller University Heilbrunn Family Center for Research Nursing awarded to RMX. Philadelphia Neurodevelopmental Cohort data used for the analyses described in this manuscript were obtained from dbGaP at http://www.ncbi.nlm.nih.gov/sites/entrez?db=gap through dbGaP accession phs000607.v3.p2.

