## Supplementary material for "Salience Network Segregation and Symptom Profiles in Psychosis Prodrome Subgroups": Supremental Information

**Supplementary Methods**

***Construction of the Neuroimaging layer***

To construct the neuroimaging layer, we computed the Pearson dissimilarity between each pair of subjects using the salience network ROIs. The Pearson dissimilarity between vectorized matrices, $\boldsymbol{q}_{\boldsymbol{1}}$ and $\boldsymbol{q}_{\boldsymbol{2}}$, is defined as the following:

$D_{P}\left( \boldsymbol{q}_{\boldsymbol{1}},\boldsymbol{q}_{\boldsymbol{2}} \right)=\frac{1-corr\left( \boldsymbol{q}_{\boldsymbol{1}},\boldsymbol{q}_{\boldsymbol{2}} \right)}{2}$,

where $\mathrm{corr}\left( \boldsymbol{q}_{\boldsymbol{1}},\boldsymbol{q}_{\boldsymbol{2}} \right)$ is the Pearson correlation coefficient between $\boldsymbol{q}_{\boldsymbol{1}}$ and $\boldsymbol{q}_{\boldsymbol{2}}$ (1). The Pearson dissimilarity ranges between 0 and 1, with higher values indicating a greater “dissimilarity” (distance) between vectors.

***Construction of the Symptom Layer***

To construct the symptom layer, we used the Mahalanobis distance, a measure of how far a single point lies from a multivariate normal distribution. If we assume the data follows a multivariate normal distribution, the Mahalanobis distance can be modified to measure how far two points lie in a multidimensional space. As such, the Mahalanobis distance between column vectors, $\boldsymbol{q}_{\boldsymbol{1}}$ and $\boldsymbol{q}_{\boldsymbol{2}}$, is defined as the following:

$D_{M}\left( \boldsymbol{q}_{\boldsymbol{1}},\boldsymbol{q}_{\boldsymbol{2}} \right)=\sqrt{\left( \boldsymbol{q}_{\boldsymbol{1}}-\boldsymbol{q}_{\boldsymbol{2}} \right)^{T}\boldsymbol{S}^{-1}\left( \boldsymbol{q}_{\boldsymbol{1}}-\boldsymbol{q}_{\boldsymbol{2}} \right)}$,

where $\boldsymbol{S}^{-1}$ is the inverse of the covariance matrix between $\boldsymbol{q}_{\boldsymbol{1}}$ and $\boldsymbol{q}_{\boldsymbol{2}}$. When computing pairwise similarity distances, $\boldsymbol{q}_{\boldsymbol{1}}$ and $\boldsymbol{q}_{\boldsymbol{2}}$ are vectors of prodromal psychosis response values for separate subjects. The Mahalanobis distance is constrained to take on positive values.

***Multiplex SBM formulation***

Let us assume we observe $n$ nodes (vertices) with *K* types of edges (links or connections) between each pair of nodes. In our study, each node is a single participant. We can represent this with a set of $K$ undirected graphs (network layers), $\left\{ X^{1},\ldots,X^{K} \right\}$, relying on the same set of $n$ nodes, $V=\left\{ 1,\ldots,n \right\}$. The nodes can be partitioned into $Q$ blocks (groups or clusters), represented by latent variables, $\left( Z_{i} \right)_{i=1,\ldots,n}$, such that $Z_{i}=q$ if node $i$ belongs to block $q\in Q$. We can define a distribution on the edge values between a given pair of nodes, $i\in V$ and $j\in V$, in graph $\left( X^{K} \right)_{k\in K}$:

$\boldsymbol{X}_{ij}^{k}|Z_{i}=q,Z_{j}=r \sim_{\mathrm{ind}} \mathcal{F}_{k}\left( \cdot;\theta_{qr}^{k} \right)$,

where $q\in Q$ and $r\in Q$ are the blocks to which $i$ and $j$ belong, respectively, and $\mathcal{F}_{k}$ is a probability distribution based on the data (2). As such, the edge values between any two nodes are driven by their block memberships. We assume that the edges are weighted with real values and that $\mathcal{F}_{k}$ is a Gaussian distribution parameterized by the following:

$\theta_{qr}^{k}=\left( \mu_{qr},\sigma_{qr}^{2} \right)$,

where $\mu_{qr}$ and $\sigma_{qr}^{2}$ are the mean and variance, respectively, of the edge weights between block $q$ nodes and block $r$ nodes. This model assumes that the $K$ edge weights between nodes $i$ and $j$ are conditionally independent to the clustering.

Furthermore, a given node has a probability associated with belonging to each block:

$\forall_{q\in Q}P\left( Z_{i}=q \right)=\alpha_{q}$,

where $\alpha_{q}$ is the probability that node $i$ belongs to block $q$ (2). Block assignment was determined by the highest value of $\alpha$ for each node. If two or more blocks had the same highest $\alpha$ for a given node, the node was randomly assigned to one of the blocks.

***Multiplex SBM estimation***

The SBM parameters were estimated using a variational expectation-maximization (EM) algorithm, and the clustering was initialized by fitting separate simple SBMs to each layer (3). Subjects were assigned to blocks using the posterior probabilities ($\alpha$ values) from the final iteration of the variational EM algorithm. We selected the optimal number of blocks (*Q*) using the integrated completed likelihood (ICL), a penalized likelihood criterion. We evaluated values of *Q* ranging from one to seven and selected the models that converged with the highest ICL above a tolerance threshold of 0.01% of the next highest value. A visualization of the optimal *Q* selection process is presented in Figure 1B, and a table of parameter estimates is presented in Figure 1C.

**Supplemental Tables**

Table S1. Psychopathology Symptom Score Statistics and Two-tailed t-test Results for the Final Sample (N=922) and the Youth (n=637) and Early Adults (n=295) Subgroups

| Psych.  Domain |  | Sample Subgroup | | Total | BH-adj.  *t*-prob. |
| --- | --- | --- | --- | --- | --- |
|  |  | Youth | Early Adults |  |  |
| ADD | Mean (S.D.) | 3.04 (2.76) | 2.20 (2.55) | 2.78 (2.72) | ***7.05×10^–5^*** |
| AGR |  | 0.89 (2.04) | 0.39 (1.04) | 0.73 (1.81) | ***3.97×10^–4^*** |
| CDD |  | 0.60 (1.15) | 0.80 (1.69) | 0.66 (1.35) | 5.16×10^–2^ |
| DEP |  | 1.25 (1.48) | 1.38 (1.52) | 1.29 (1.49) | 2.62×10^–1^ |
| GAD |  | 0.86 (1.16) | 0.88 (1.04) | 0.86 (1.12) | 7.69×10^–1^ |
| MAN |  | 1.78 (2.41) | 1.47 (2.11) | 1.69 (2.33) | 6.88×10^–2^ |
| OCD |  | 1.63 (2.56) | 0.99 (1.86) | 1.43 (2.39) | ***5.29×10^–4^*** |
| ODD |  | 1.64 (1.71) | 1.40 (1.48) | 1.57 (1.64) | 5.16×10^–2^ |
| PAN |  | 0.47 (1.05) | 0.52 (0.88) | 0.48 (1.00) | 5.05×10^–1^ |
| PHB |  | 2.33 (2.40) | 1.91 (2.03) | 2.20 (2.30) | ***1.83×10^–2^*** |
| PSY |  | 0.80 (1.41) | 0.34 (0.88) | 0.66 (1.29) | ***7.22×10^–6^*** |
| PTD |  | 1.01 (1.38) | 1.38 (1.87) | 1.13 (1.55) | ***2.43×10^–3^*** |
| SEP |  | 1.22 (1.68) | 0.86 (1.23) | 1.11 (1.56) | ***2.43×10^–3^*** |
| SOC |  | 1.72 (1.69) | 1.37 (1.65) | 1.61 (1.69) | ***7.70×10^–3^*** |
| SUI |  | 0.27 (0.54) | 0.38 (0.66) | 0.30 (0.58) | ***1.42×10^–2^*** |
| SIP ^a^ |  | 0.89 (1.09) | 0.55 (0.84) | 0.78 (1.03) | ***3.27×10^–5^*** |

ADD, attention deficit hyperactive disorder; AGR, agoraphobia; CDD, conduct disorder; DEP, depression; GAD, generalized anxiety disorder; MAN, mania; OCD, obsessive compulsive disorder; ODD, oppositional defiant disorder; PAN, panic disorder; PHB, specific phobias; PSY, psychosis spectrum; PTD, post traumatic disorder; SEP, separation anxiety disorder; SOC, social anxiety disorder; SUI, suicide; SIP, psychosis prodrome.

Significant two-tailed *t*-test results (BH-adj. *t*-prob.) are bolded and italicized. Significance was determined based on a threshold of 0.05. The Benjamini-Hochberg method was used to adjust for multiple comparisons.

a) For the ordinal psychosis prodrome (SIP) items, the average response value per subject was computed. For each psychopathology domain with binary response items (i.e. all psychopathology domains except for SIP), the number of positive responses per subject was computed.

Table S2*.* Psychopathology Symptom Score Statistics and Two-tailed *t*-test Results for Each Block among the Youth (*n*=637; Block 1: *n*=458; Block 2: *n*=179)

| Psych.  Domain | Block | | BH-adj. *t*-prob. |
| --- | --- | --- | --- |
|  | 1 | 2 |  |
|  | Mean (S.D.) | |  |
| ADD | 3.54 (2.78) | 1.76 (2.24) | ***5.47×10^–13^*** |
| AGR | 0.98 (1.94) | 0.65 (2.27) | 7.32×10^–2^ |
| CDD | 0.71 (1.24) | 0.30 (0.80) | ***4.63×10^–5^*** |
| DEP | 1.48 (1.53) | 0.69 (1.16) | ***2.35×10^–9^*** |
| GAD | 0.94 (1.20) | 0.64 (1.03) | ***3.74×10^–3^*** |
| MAN | 2.18 (2.53) | 0.77 (1.71) | ***6.54×10^–11^*** |
| OCD | 2.03 (2.81) | 0.59 (1.29) | ***3.06×10^–10^*** |
| ODD | 1.90 (1.77) | 0.99 (1.34) | ***2.96×10^–9^*** |
| PAN | 0.59 (1.18) | 0.14 (0.42) | ***1.43×10^–6^*** |
| PHB | 2.63 (2.56) | 1.54 (1.73) | ***3.99×10^–7^*** |
| PSY | 1.04 (1.52) | 0.20 (0.80) | ***4.65×10^–11^*** |
| PTD | 1.13 (1.45) | 0.72 (1.13) | ***6.60×10^–4^*** |
| SEP | 1.37 (1.76) | 0.86 (1.37) | ***6.60×10^–4^*** |
| SOC | 1.91 (1.71) | 1.24 (1.56) | ***1.13×10^–5^*** |
| SUI | 0.32 (0.59) | 0.12 (0.36) | ***2.05×10^–5^*** |
| SIP ^a^ | 1.24 (1.10) | 0.00 (0.00) | ⸻ |

ADD, attention deficit hyperactive disorder; AGR, agoraphobia; CDD, conduct disorder; DEP, depression; GAD, generalized anxiety disorder; MAN, mania; OCD, obsessive compulsive disorder; ODD, oppositional defiant disorder; PAN, panic disorder; PHB, specific phobias; PSY, psychosis spectrum; PTD, post traumatic disorder; SEP, separation anxiety disorder; SOC, social anxiety disorder; SUI, suicide; SIP, psychosis prodrome.

Significant two-tailed *t*-test results (BH-adj. *t*-prob.) are bolded and italicized. Significance was determined based on a threshold of 0.05. The Benjamini-Hochberg method was used to adjust for multiple comparisons.

**a) For the ordinal psychosis prodrome (SIP) items, the average response value per subject was computed. For each psychopathology domain with binary response items (i.e. all psychopathology domains except for SIP), the positive (“yes”) response count per subject was computed. Since the multiplex SBM was fit to the SIP response data, cross validation was conducted using response data from all psychopathology domains except SIP. Therefore, there are no *t*-test results for the SIP domain.**

Table S3. Psychopathology Symptom Score Statistics and Two-tailed *t*-test Results for Each Block among the Early Adults (*n*=285; Block 1: *n*=173; Block 2: *n*=112)

| Psych.  Domain | Block | | BH-adj. *t*-prob. |
| --- | --- | --- | --- |
|  | 1 | 2 |  |
|  | Mean (S.D.) | |  |
| ADD | 2.74 (2.67) | 1.37 (2.12) | ***2.06×10^–5^*** |
| AGR | 0.60 (1.28) | 0.06 (0.24) | ***4.01×10^–5^*** |
| CDD | 1.00 (1.96) | 0.48 (1.10) | ***1.14×10^–2^*** |
| DEP | 1.64 (1.55) | 0.98 (1.40) | ***5.09×10^–4^*** |
| GAD | 1.04 (1.12) | 0.63 (0.83) | ***1.26×10^–3^*** |
| MAN | 1.93 (2.30) | 0.75 (1.52) | ***1.40×10^–5^*** |
| OCD | 1.38 (2.21) | 0.38 (0.84) | ***2.06×10^–5^*** |
| ODD | 1.69 (1.54) | 0.96 (1.27) | ***7.45×10^–5^*** |
| PAN | 0.67 (0.95) | 0.29 (0.70) | ***5.15×10^–4^*** |
| PHB | 2.24 (2.20) | 1.40 (1.61) | ***7.53×10^–4^*** |
| PSY | 0.53 (1.08) | 0.05 (0.23) | ***2.06×10^–5^*** |
| PTD | 1.70 (2.04) | 0.88 (1.44) | ***4.26×10^–4^*** |
| SEP | 1.06 (1.22) | 0.55 (1.19) | ***8.20×10^–4^*** |
| SOC | 1.59 (1.76) | 1.04 (1.42) | ***5.92×10^–3^*** |
| SUI | 0.53 (0.74) | 0.14 (0.42) | ***9.78×10^–6^*** |
| SIP ^a^ | 0.91 (0.92) | 0.00 (0.00) | ⸻ |

ADD, attention deficit hyperactive disorder; AGR, agoraphobia; CDD, conduct disorder; DEP, depression; GAD, generalized anxiety disorder; MAN, mania; OCD, obsessive compulsive disorder; ODD, oppositional defiant disorder; PAN, panic disorder; PHB, specific phobias; PSY, psychosis spectrum; PTD, post traumatic disorder; SEP, separation anxiety disorder; SOC, social anxiety disorder; SUI, suicide; SIP, psychosis prodrome.

Significant two-tailed *t*-test results (BH-adj. *t*-prob.) are bolded and italicized. Significance was determined based on a threshold of 0.05. The Benjamini-Hochberg method was used to adjust for multiple comparisons.

**a) For the ordinal psychosis prodrome (SIP) items, the average response value per subject was computed. For each psychopathology domain with binary response items (i.e. all psychopathology domains except for SIP), the positive (“yes”) response count per subject was computed. Since the multiplex SBM was fit to the SIP response data, cross validation was conducted using response data from all psychopathology domains except SIP. Therefore, there are no *t*-test results for the SIP domain.**

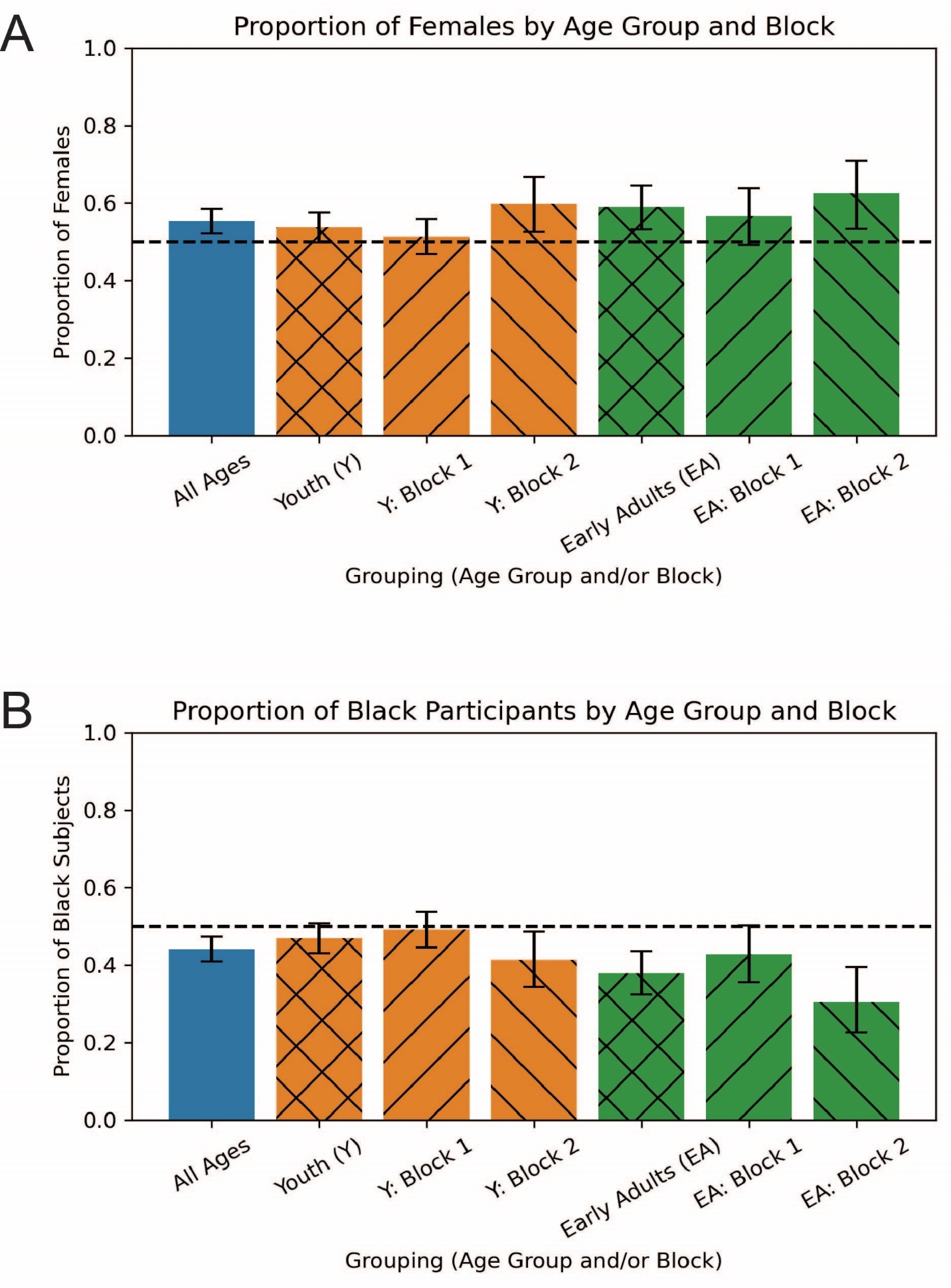

Figure S1. Visualization of participant demographics broken down by age group (*N*=922; youth [Y]: *n*=637; early adults [EA]: *n*=285) and block (Y Block 1: *n*=458; Y Block 2: *n*=179; EA Block 1: *n*=173; EA Block 2: *n*=112).

A bar plot with 95% confidence intervals for the proportion of female participants in each grouping is shown in (**A**). Group comparisons of the gender distributions between age groups and blocks were significant ($\chi^{2}$=15.96, *p*=1.15×10^–3^), but post hoc comparisons between blocks were not (youth: $\chi^{2}$=3.38, *p*=6.61×10^–2^; early adults: $\chi^{2}$=0.74, *p*=3.91×10^–1^).

A bar plot with 95% confidence intervals for the proportion of Black participants in each grouping is shown in (**B**). While the race demographic contains four categories (Black, White, mixed race, and “other”), most participants identify as either Black or White. Thus, race can be presented as a single measure (i.e., the proportion of Black subjects) without much loss of information. Group comparisons of the race distributions between age groups and blocks were significant ($\chi^{2}$=19.73, *p*=6.19×10^–3^), but post hoc comparisons between blocks were not (youth: $\chi^{2}$=3.26, *p*=3.53×10^–1^; early adults: $\chi^{2}$=4.75, *p*=1.91×10^–1^).

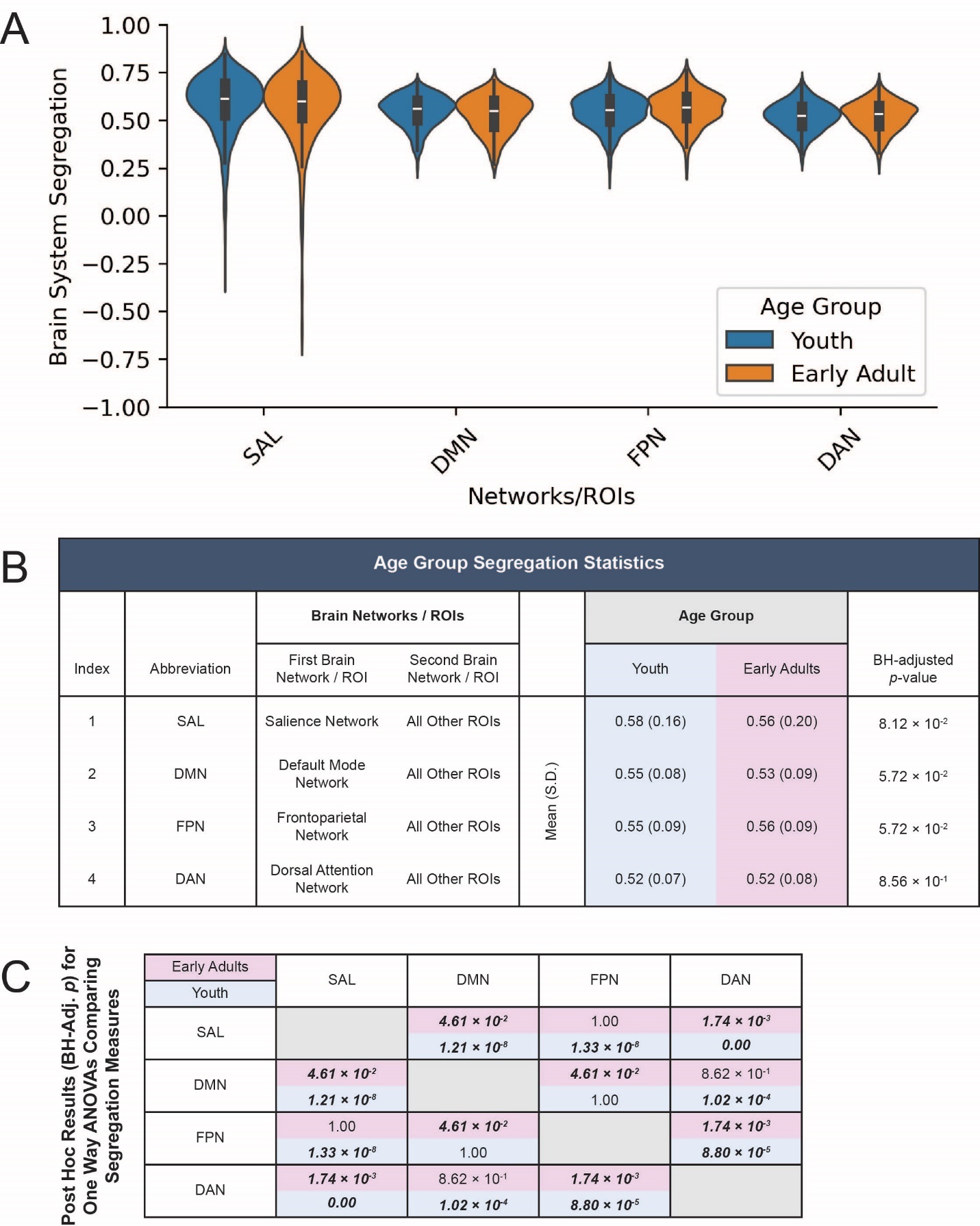

Figure S2. Brain system segregation values for the youth (*n*=637) and early adult (*n*=285) participants prior to subgroup identification (community detection).

A visualization of the distributions of segregation values broken down by age group is shown in (**A**).

A table with segregation statistics (means and standard deviations) for the two age groups and two-tailed *t*-test results (BH-adjusted *p*-value) for comparisons of means between the groups is shown in (**B**). None of the *t*-test comparisons were significant.

A symmetric matrix with post-hoc comparison (two-tailed *t*-test) results for one-way ANOVAs that assess differences between the four segregation measures is shown in (**C**). An ANOVA was conducted separately for the youth (*F*=39.40, *p*=6.95×10^–25^) and the early adults (*F*=7.35, *p*=7.00×10^–5^). Significant pair-wise *t*-test results (BH-adj. *p*) are bolded and italicized.

Significance was determined based on a threshold of 0.05. The Benjamini-Hochberg method was used to adjust for multiple comparisons.

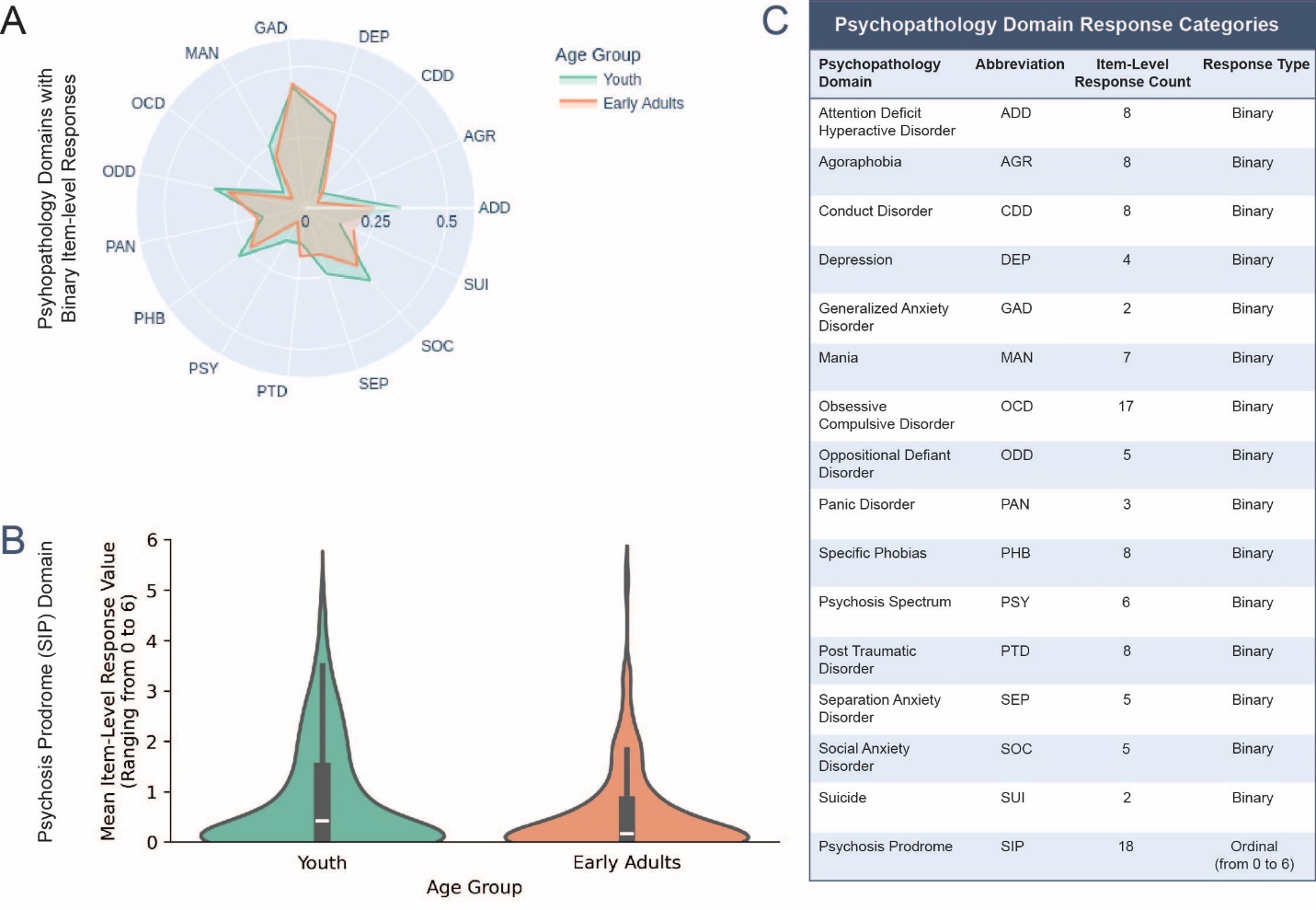

Figure S3. Psychopathology symptom scores for the youth (*n*=637) and early adult (*n*=285) participants prior to subgroup identification (community detection).

The mean positive (“yes”) response count per subject for each of the 15 non-prodromal psychopathology domains is shown in (**A**). The means are scaled to range from 0 to 1 to better visualize the differences between blocks.

The distributions of the mean response values for the prodromal psychosis items are shown in (**B**).

A table displaying the abbreviation, number of item-level responses, and response value type (binary or ordinal) for each psychopathology domain is shown in (**C**).
